# Compromised External Validity: Federally Produced *Cannabis* Does Not Reflect Legal Markets

**DOI:** 10.1101/083444

**Authors:** Daniela Vergara, L. Cinnamon Bidwell, Reggie Gaudino, Anthony Torres, Gary Du, Travis C. Ruthenburg, Kymron deCesare, Donald P. Land, Kent E. Hutchison, Nolan C. Kane

## Abstract

As the most widely used illicit drug, the basis of the fastest growing major industry in the US, and as a source of numerous under-studied psychoactive compounds, understanding the psychological and physiological effects of *Cannabis* is essential. The National Institute on Drug Abuse (NIDA) is designated as the sole legal producer of *Cannabis* for use in US research studies. We sought to compare the chemical profiles of *Cannabis* varieties that are available to consumers in states that have state-legalized use *versus* what is available to researchers interested in studying the plant and its effects. Our results demonstrate that the federally produced *Cannabis* has significantly less variety and lower concentrations of cannabinoids. Current research, which has focused on material that is far less diverse and less potent than that used by the public, limits our understanding of the plant’s chemical, biological, psychological, medical, and pharmacological properties. Investigation is urgently needed on the diverse forms of *Cannabis* used by the public in state-legal markets.

## Introduction

The United States has witnessed enormous changes concerning public acceptance of marijuana. Use has more than doubled since 2002, across all genders, ethnicities and socioeconomic status^1^. Considering changes on the cultural, political, and legal fronts, research on the effects of *Cannabis* products that are consumed though legal outlets in states that have legalized is urgently needed.

The *Cannabis* plant is unique in producing a diversity of cannabinoids, a terpenoid chemical compound that interacts with the endocannabinoid system in the brain and nervous system^2^. One of the primary cannabinoids produced, Δ-9-tetrahydrocannabinolic acid (THCA), is converted to the neutral form Δ-9-tetrahydrocannabinol (THC) when heated, e.g. by smoking. THC interacts with the endocannabinoid system producing a wide range of physiological and neurological effects. Studies have found that marijuana’s effects on mood, reward, and cognitive dysfunction appear to follow a dose dependent function based on the THC content^3^. Due to this and other purported psychotropic effects, THCA has been actively selected for by the *Cannabis* industry^4^, and varieties containing more than 30% THCA by weight have been cultivated^5^. In addition to THC, marijuana’s effects are likely related to a number of other compounds^6,7^, including nearly 74 different cannabinoids^8^ present at varying ratios across strains. For example, cannabidiolic acid (CBDA), another cannabinoid produced by the plant, is converted to cannabidiol (CBD) when heated. CBD may mitigate the effects of THC and may have other beneficial effects^9–18^. Demand for high CBDA plants has increased, due to potential therapeutic uses for cancer^19^ and epilepsy^20,21^. Other important cannabinoids produced by the plant include cannabigerol (CBG)^22^, cannabichrome (CBC)^23^, and Δ-9-tetrahydocannabivarin (THCV)^24^.

Because research universities across the nation have national grants and must verify compliance with federal law, scientists at these institutions are restricted to research with the only federally legal source of *Cannabis*. Our current understanding of the effects of marijuana in humans (e.g. on mood, cognition, or pain) has therefore relied exclusively on government-grown marijuana, often administered in a laboratory setting^25–27^. Thus, nearly all published US laboratory studies have used *Cannabis* material obtained from the National Institutes of Health/National Institute on Drug Abuse (NIDA) supply, the only federally legal source for *Cannabis* plant material. At the same time, dispensary-grade *Cannabis* available to consumers in state-regulated markets is becoming increasingly potent and diverse. Strains differ substantially in potency and cannabinoid content, and hence, are likely to differ in terms of their effects^4^. Strains bred for high THCA content are thought to result in greater levels of intoxication as well as differing psychological and physiological effects compared to strains bred for high CBDA content. Accordingly, NIDA has recently developed plant material with varying levels of cannabinoids for research purposes, but the extent to which government *Cannabis* is consistent with *Cannabis* produced in the private market is not clear.

To address the critical question of whether the potency and variety of NIDA-provided *Cannabis* reflects products available to consumers through state-regulated markets, we compared the cannabinoid variation and potency from plants from four different cities in the US where *Cannabis* is legal for medical or recreational reasons (Denver, Oakland, Sacramento, and Seattle; cannabinoid data provided by Steep Hill Labs Inc.) to the cannabinoid content of plants supplied for research purposes by NIDA, using the data publicly available on their website ^28^. Table 1 shows the sample sizes for all locations.

**Table 1.** Sample sizes. Sample sizes for each cannabinoid at the different locations. Denver and Seattle lack samples for CBN and CBC. The first column represents the total sample sizes for each cannabinoid, the >1 column represents the number of samples that produced more than 1% content of each cannabinoid. No location had strains that produced >1% CBN or CBC. The last column represents the sample sizes used for the PCA, which was performed only with samples from Oakland, Sacramento, and NIDA. The last row represents the samples that produce more than 1% for both THC and CBD.

|  | Denver |  | NIDA |  |  | Oakland |  |  | Sacramento |  |  | Seattle |  |
| --- | --- | --- | --- | --- | --- | --- | --- | --- | --- | --- | --- | --- | --- |
|  | TOTAL | >1 | TOTAL | >1 | PCA | TOTAL | >1 | PCA | TOTAL | >1 | PCA | TOTAL | >1 |
| **CBD** | 1141 | 42 | 98 | 56 | 90 | 755 | 110 | 481 | 981 | 101 | 981 | 103 | 4 |
| **CBN** | - | - | 98 | - | 90 | 481 | - | 481 | 981 | - | 981 | - | - |
| **THC** | 1141 | 1112 | 98 | 64 | 90 | 755 | 692 | 481 | 981 | 952 | 981 | 103 | 103 |
| **CBG** | 992 | 98 | 96 | 1 | 90 | 481 | 41 | 481 | 981 | 259 | 981 | 103 | 12 |
| **THC-V** | 992 | 12 | 96 | - | 90 | 481 | 6 | 481 | 981 | 21 | 981 | 103 | 1 |
| **CBC** | - | - | 96 | - | 90 | 481 | - | 481 | 981 | 2 | 981 | - | - |
| **THC & CBD >1** | 21 | - | 24 | - | - | 77 | - | - | 81 | - | - | 4 | - |

## Results

NIDA differs from all other locations except Seattle in production of CBD (Fig. 1A), and differs significantly from all other locations in production of THC. NIDA has the lowest CBD and THC percent with a mean and s.d. of 6.16 ± 2.43%, and 5.15 ± 2.60% respectively. Sacramento has the highest percent CBD with 12.83 ± 4.73% and Seattle has the highest percent THC with 19.04 ± 4.43%. There are significant differences between the percent of both CBD and THC between US city locations, in addition to differences with NIDA (Fig. 1A).

**Fig. 1.**
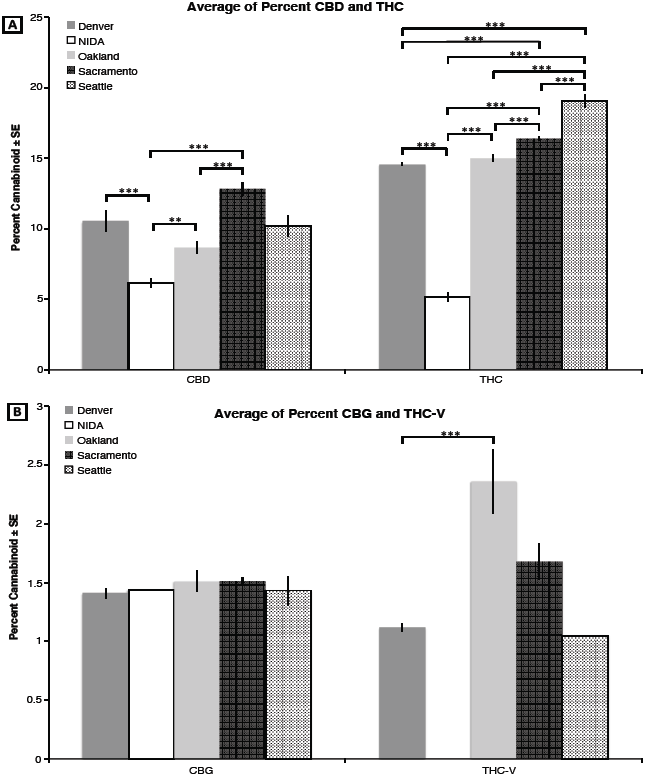
Average percent cannabinoids for five different locations. A. CBD (N=313) and THC (N=2923). **B.** CBG (N=411) and THC-V (40). Significant values between the comparisons are given in the horizontal bars above: *** P<0.001; ** P <0.01; and * P<0.05.

CBG production does not differ in any location. *Cannabis* plants grown in all locations produce very little CBG, particularly NIDA with only a single sample producing more than 1% CBG (Fig. 1B). THC-V is also produced in low quantities in all locations. The only statistically significant difference is between Denver, whose mean and s.d is 1.12 ± 0.13%, and Oakland 2.35 ± 0.68% (P<0.001; fig. 1B). Importantly, Seattle has only one (1) sample and NIDA lacks any plants that produce more than 1% THC-V.

Two analyses were used to examine the phytochemical diversity found in each location. In the first analysis, the variability and range of each cannabinoid was calculated (figure 2). This analysis shows that for three of the four cannabinoids, NIDA has the lowest variability. In addition, the potencies of THC and CBD across sites (figure 3) suggest a greater diversity in both potency and ratio in the private market. In other words, the federal strains show limited diversity in the total cannabinoid levels, in the cannabinoids that are present, and in the ratio of cannabinoids (figure S1).

**Fig. 2.**
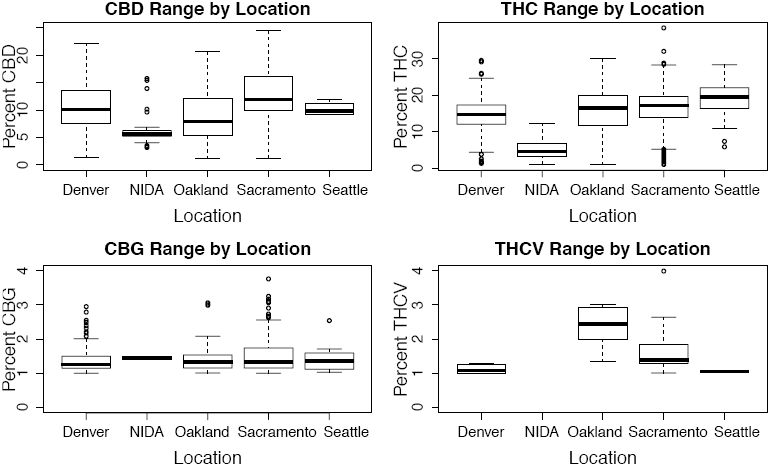
Median and range for cannabinoids by location. Median (line within the box), 25^th^ and 75^th^ percentile (bottom and top of the box respectively), and range (bars outside the box).Outliers are dots outside the box and range. The Y axis differs by panel.

**Fig. 3.**
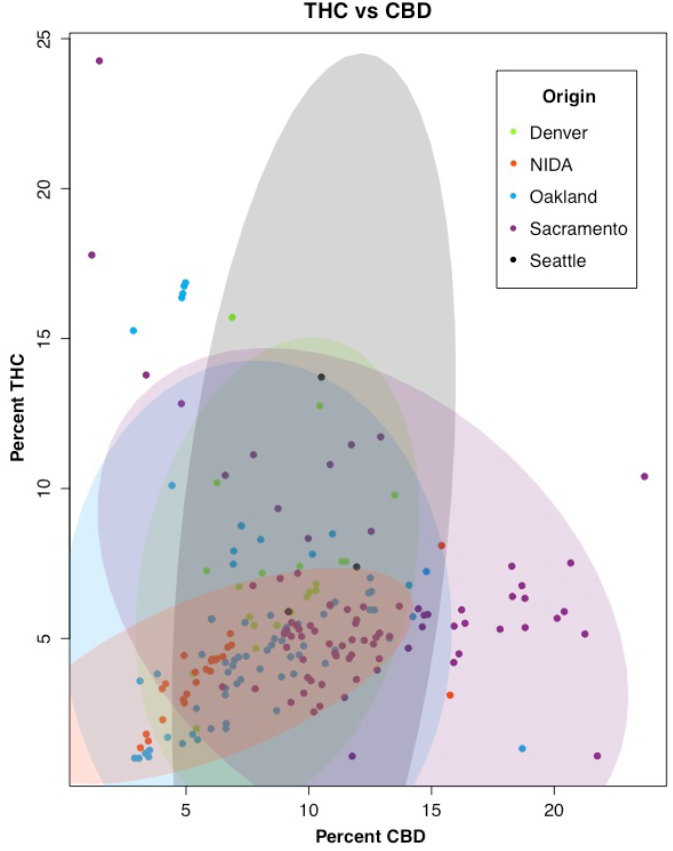
The diversity and variability of *Cannabis* samples across sites in terms of THC, CBD. The ellipses represent 95% confidence (N=1152).

PCA (fig. 4) shows that 53.1% of the overall cannabinoid variation is explained by PC1 and PC2, with PC1 explaining 30.8% and PC2 explaining 22.3%. PCA shows that the overall cannabinoid content in Oakland and Sacramento is very similar, since the points overlap with each other, even though Sacramento has more variation. Most of NIDA’s samples cluster within the ones from Sacramento and Oakland. Additionally, NIDA’s 95% confidence ellipse mostly lies within the Sacramento and Oakland ellipses. In other words, the variation in cannabinoids from NIDA can be found in Oakland and Sacramento. However, the variation from Sacramento and Oakland is not captured by the NIDA strains. Therefore across all cannabinoids, the government source of *Cannabis* is limited in diversity, not reflecting the one widely available to consumers in state markets. Additionally we established with the k-means cluster analysis that the best number of clusters given the data from the PCA analysis was two. These two groups are clearly portrayed in the PCA graph with PC1 against PC2 (figs. 4 and S2), revealing that NIDA’s samples are present only in one of the clusters. Therefore, the cannabinoid diversity from the private market is represented in both clusters, while that from the federal cannabinoids is almost entirely found only in one of the clusters, demonstrating again their lack of variation.

**Fig. 4.**
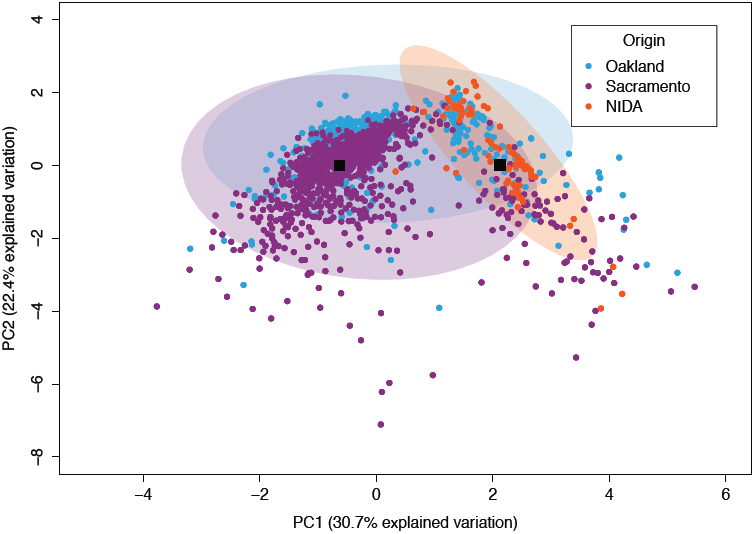
PC1 vs PC2 for three locations. Most of the points from the two main PC axes overlap demonstrating similarities between the three locations in their content. The black boxes represent the means of the two clusters after the k-means analysis.

## Discussion

The objective of this research was to determine whether *Cannabis* produced by the government for research reflects the *Cannabis* that is widely available in state regulated markets. The data demonstrate that *Cannabis* plants currently grown for NIDA are not representative of plants consumed by recreational and medicinal users through state-legalized markets across the nation. *Cannabis* flower available from dispensaries appears to be more potent and diverse in their cannabinoid content.

This underrepresentation in the NIDA’s cultivars is problematic for investigation in several areas, including chemistry, biochemistry, genomics, biology, psychology, but particularly for medical research. These data suggest that the NIDA strains underrepresent the genomic variation of cultivars with higher cannabinoid levels and the genomic variation that is found in state-legalized markets ^29,30^. Medical research using only a limited number of varieties can be misleading, because variation in the amounts and ratios of cannabinoids may have a significant impact on the outcomes of the studies. Particularly as cannabinoids can have dramatic opposing effects and complex interactions with each other^9–18^, investigations that only use one source of material may hinder our understanding of pharmacological and therapeutic effects of *Cannabis*.

Studies reporting on effects of *Cannabis* using NIDA strains will continue to lack external validity, possibly underestimating the effects of more potent strains that are widely available. Compounding this problem, is the fact that the public availability of high-potency *Cannabis* has increased in recent years^1^. Given our data and recent reviews that have suggested that the greater potency of today’s marijuana, compared to earlier decades^4^, may lead to significantly greater levels of intoxication and harm, it is important for research to begin understanding consequences and impact of using the publicly available *Cannabis.* The knowledge gap between what we know from studies using government grown *Cannabis* and ***what we should know*** about the effects of *Cannabis* in the real world will continue to widen with the progressive decriminalization and accessibility of high-potency, dispensary-grade *Cannabis*. This problem can only be addressed by establishing legal methods for the nation’s scientists to access the *Cannabis* more similar to what is sold and consumed in state-regulated marketplaces for research on the potential for harm and medical applications.

Despite this being the most complete cannabinoid analysis to date, it has a number of limitations. First, data analyzed in this study were collected by separate facilities, NIDA and Steep Hill, which may introduce biases. Inter-laboratory comparative analysis between different methods of testing *Cannabis* products (e.g., different equipment used for this comparison and the various facilities that offer chemotype testing) has been limited. This limitation is largely driven by federal laws that prevent third parties (e.g., universities) from conducting such studies. Lastly, Steep Hill data only includes strains tested at their locations and are not necessarily representative of all *Cannabis* available to consumers. However, with 2980 samples tested, common varieties are well represented. Similarly, our analysis includes the potential current pool of strains listed as available by the government for research purposes; however the *Cannabis* varieties produced historically by NIDA (and used in NIDA-funded studies published prior to 2012, when these additional NIDA strains became available) are far less potent. Thus, while our analysis focuses on currently available strains, the discrepancy between publicly-available marijuana and that used in most existing research is even greater than what we report here.

Moreover, this analysis is limited to six cannabinoids reported for the NIDA strains, while additional compounds are known to be important^2,7,8^. Chemical analyses of *Cannabis* in commercial testing labs include numerous cannabinoids and terpenoids, which vary between lineages^31^ and have important physiological effects^2,7,8^. Compounds not reported for the NIDA strains may represent additional important differences between the federally approved strains and widely used material. With the reported cannabinoids in this study, it seems unlikely that NIDA strains will ever represent the broader marketplace. However, it is worth noting that the strains available through NIDA may compare favorably to individual dispensaries in terms of diversity of cannabinoid levels and ratios of THC to CBD.

Additionally, *Cannabis* flower is one form of *Cannabis* available in state regulated markets, with concentrates and edibles also widely used. It is critical to note that as of May 2016 on the federal website, there is no source of concentrates or edibles for research. Therefore, there is almost no research on the effects of cannabinoids in extract or edible form, even though in Colorado alone, approximately 650,000 edible units are sold each month. Given the diversity of the products, the federal government is unlikely to be able to produce *Cannabis* in a way that reflects the diversity products used by the public in states where it is legal.

In conclusion, this study offers a comparison between six cannabinoids from *Cannabis* produced in various cities in the US and the NIDA supply farm. The data suggest that *Cannabis* produced by NIDA is less diverse in variety and less potent in the amount of cannabinoids. Because doing federally approved research requires the use of government produced *Cannabis*, the current situation is a significant impediment to research that seeks to clarify the potential harms or benefits. In recent years, federal sources have pursued diversification of their strains with a goal of increasing the diversity and potency of research *Cannabis*. The research presented here provides concrete data that can inform these changes, so that *Cannabis* available to researchers in the future can better reflect the types of products widely-used by the public.

## Methods

Cannabinoids from multiple strains in four cities of the US Denver, Oakland, Sacramento, and Seattle were measured by Steep Hill Labs, Inc. These samples were not randomly chosen for two main reasons: First, because we rely on the locations where Steep Hill has facilities, and second, because even though dispensary owners and *Cannabis* producers are required by law to test their product in some of those jurisdictions, it is a choice to select between the multiple companies that provide these services. However, Steep Hill is the only company that tests for 17 cannabinoids and ten terpenes^32^, and has multiple facilities in cities where *Cannabis* is legal, medically and/or recreationally. The use of the same testing procedures across multiple marketplaces allows us to compare cannabinoid levels among the largest state markets. Moreover, this dataset includes many widely used strains, as well as minor ones.

Cannabinoid measurements are performed on the flower of female plants, where most of the cannabinoids are produced^33, 34^. Steep Hill samples were weighted and their phytochemicals extracted to then be filtered and diluted. All Steep Hill samples were measured using liquid chromatography (LC), in Denver with Agilent LC equipment, in Seattle and Sacramento using Shimadzu LC equipment, and in Oakland using both types of machinery. The data from NIDA was obtained from their website on November 15, 2015^28^. Details about data collection or the equipment used was not currently specified. Total sample sizes for each of the cannabinoids by location are given in Table 1. Even though Steep Hill measures additional cannabinoids, our analyses focused on cannabinoids shared between the NIDA and SteepHill datasets (N=6). NIDA uses gas chromatography for their analysis ^35–37^, which only measures the neutral form of the cannabinoids. Thus, we transformed the acidic form of each cannabinoid by multiplying each LC value by the ratio of molecular masses of the neutral cannabinoid to the acidic cannabinoid, which represents the mass ratio of the neutral relative to acidic forms after decarboxylation. The state mandated value for this conversion is approximately 0.88 and several states such as Washington^38^ and Nevada^39^ mandate reporting of total THC from HPLC analysis using this calculation. We then added the converted values to the neutral form in our measurements, the result is equivalent to the measurements taken by NIDA. We also performed a separate analysis using the conversion factor reported by Dussy and collaborators of 0.68^40^. Even though both datasets using the two conversion rates differ from each other, the overall results and conclusions are the same: a significantly lower total diversity and total level of cannabinoids from the NIDA samples. Therefore, we are only presenting the sample sizes and results using the state-mandated conversion rate of 0.88.

In order to analyze the cannabinoid potency information across the sites, we first selected only those tests that demonstrated 1% or greater for the specific cannabinoid under analysis. This method allowed us to more accurately report the concentration of particular cannabinoids across strains, many of which are bred for high production of a specific cannabinoid and low concentrations of another particular cannabinoid. Due to the absence of samples that produced more than 1% CBN and CBC, we excluded these two cannabinoids from the analysis. An ANOVA was then performed for each cannabinoid with location as a factor and a posterior posthoc analysis using Tukey, except for THC-V where we used Bonferroni (fig. 1). We determined the cannabinoid range on a box and whiskers plot (fig. 2) to visualize the array, median, minimum, and maximum of each compound by location. Additionally, we determined which samples produced more than 1% in both THC and CBD (fig. 3), indicating functional copies of both THCA and CBDA-synthases, and calculated the ratio (THC/CBD), and with this ratio performed four F-tests comparing NIDA to the other four locations. Finally, to further understand the variation between locations in their overall cannabinoid composition, we performed Principal Components Analysis (PCA; fig. 4) with the Car package^41^ from R statistical software. For this analysis all samples, including the ones that produced less than 1% cannabinoids were used, however, both Denver and Seattle were excluded from the PCA due to the absence of CBN and CBC. The total sample size for this PCA is given in Table 1. We used the same dataset from the PCA to calculate k-means clustering to understand the number of partitions and their means given our data. We used the R statistical framework to perform all analyses. All code is available on www.github.com/KaneLab.

## Acknowledgments

We thank S. Tittes and K. Keepers for comments, bioinformatics help, and suggestions. This research was supported by donations to the University of Colorado Foundation gift fund 13401977-Fin8 to NCK, by donations to the Agricultural Genomics Foundation, and by K23DA033302 to LCB. The raw data for this research will be submitted to Dryad upon acceptance, and also to the website www.agriculturalgenomics.org.

## Author contributions

DV analyzed the data and wrote the first draft of the manuscript; LCB, NCK, KEH directed the project, contributed to statistical analysis and manuscript revisions; RG, DPL, TCR, KdC designed and supervised data collecting methods; AT, GD, TCR collected and added data to database; KEH conceived the project. All authors contributed to manuscript preparation.

